# Spinal cord injury in mice amplifies anxiety: a novel light-heat conflict test exposes increased salience of anxiety over heat

**DOI:** 10.1101/2023.01.13.523970

**Authors:** Sydney E. Lee, Emily K. Greenough, Laura K. Fonken, Andrew D. Gaudet

**Author notes:** Corresponding Author: Andrew Gaudet, Department of Psychology, University of Texas at Austin, 108 E. Dean Keeton Street, Stop A8000, Seay Building, Room 4.208, Austin, TX 78712, USA.

## Abstract

Spinal cord injury (SCI) predisposes individuals to anxiety and chronic pain. Anxiety- and pain-like behavior after SCI can be tested in rodents, yet commonly used tests assess one variable and may not replicate effects of SCI or sex differences seen in humans. Thus, novel preclinical tests should be optimized to better evaluate behaviors relating to anxiety and pain. Here, we use our newly developed conflict test – the Thermal Increments Dark-Light (TIDAL) test – to explore how SCI affects anxiety- vs. pain-like behavior, and whether sex affects post-SCI behavior. The TIDAL conflict test consists of two plates connected by a walkway; one plate remains illuminated and at an isothermic temperature, whereas the other plate is dark but is heated incrementally to aversive temperatures. Control mice are tested with both plates illuminated (thermal place preference). Female and male mice received moderate T9 contusion SCI or remained uninjured. At 7 days post-operative (dpo), mice with SCI increased dark plate preference throughout the TIDAL conflict test compared to uninjured mice. SCI increased dark plate preference for both sexes, although female (vs. male) mice remained on the heated-dark plate to higher temperatures. Mice with SCI that repeated TIDAL at 7 and 21 dpo showed reduced preference for the dark-heated plate at 21 dpo. Overall, in female and male mice, SCI enhances the salience of anxiety (vs. heat sensitivity). The TIDAL conflict test meets a need for preclinical anxiety- and pain-related tests that recapitulate the human condition; thus, future rodent behavioral studies should incorporate TIDAL or other conflict tests to help understand and treat neurologic disorders.

## Introduction

Anxiety is a common secondary condition associated with spinal cord injury (SCI). In a recent meta-analysis, 15-32% of individuals with SCI self-reported experiencing anxiety at time of assessment (Le and Dorstyn, 2016). Similarly, anxiety-induced behavior may be increased after injury in mice: mice with SCI show decreased percent time in the open arm on the elevated plus maze and reduced percent center zone time on the open field test, indicating increased anxiety-like behavior (Fukutoku et al., 2020). However, findings on SCI-induced anxiety in rodent models are mixed and anxiety tests typically focus on single variables. Therefore, novel behavioral assays could better address how SCI modulates anxiety, and how this interacts with other stressors (e.g. pain).

Another common consequence of SCI is chronic pain, which is experienced by 65-80% of individuals with SCI (Siddall et al., 1999). Unfortunately, neuropathic pain is often intractable to existing analgesics, so there is a need to use preclinical models and tests to unveil promising therapeutic targets (Anderson, 2004; Collinger et al., 2013; Lo et al., 2016). Common rodent assays for identifying pain-like behaviors include the von Frey test (for mechanical allodynia) and the Hargreaves test (for heat hyperalgesia). These tests demonstrate that rodents with SCI exhibit mechanical and thermal hypersensitivity (Brown et al., 2021; Detloff et al., 2013; Gaudet et al., 2017; Gaudet et al., 2021; McFarlane et al., 2020). Unfortunately, these reflexive tests are overly simplified and do not fully capture the affective component of the pain experience, which limits their utility for identifying mechanisms underlying neuropathic pain (Burma et al., 2017; Kramer et al., 2017). Therefore, it is important to develop new tests that more closely model the human pain experience. One promising strategy is to incorporate a pain-eliciting stimulus in parallel with a conflicting stimulus, which could better illuminate pain-related behaviors.

Anxiety and pain often co-occur, and share partially overlapping neurocircuitry and neuroinflammatory underpinnings. Indeed, >50% of individuals with chronic pain exhibit symptoms of anxiety and/or depression (Dahan et al., 2014; Von Korff and Simon, 1996), and 45% of individuals with anxiety experience chronic pain (vs. 29% of non-anxious population (Askari et al., 2017)). In humans and rodents, neuroinflammation sensitizes brain regions involved in threat detection, learning, reward, and anxiety (e.g., amygdala; hippocampus; insula; prefrontal and anterior cingulate cortex) (Bekhbat and Neigh, 2018). Studies in rodent models have revealed neuroinflammatory mechanisms mediating anxiety-like behavior. For instance, mice exposed to social defeat stress display neuroinflammatory activation and anxiety-like behavior, and anxiety-like behaviors are reduced by ablating key pro-inflammatory signaling via IL-1ß receptor (Wohleb et al., 2014). Thus, uncovering SCI-elicited changes in anxiety- vs. pain-related behaviors could aid development of treatments for one or both conditions.

Here, we investigate how SCI affects anxiety- vs. pain-like behavior using a newly developed conflict test – the Thermal Increments Dark-Light (TIDAL) test (Lee et al., 2022). The TIDAL conflict test apparatus incorporates two temperature-controlled plates in identically sized chambers – one illuminated chamber, with a plate maintained at an isothermic temperature; and one dark chamber, which contains a plate that incrementally increases to aversive temperatures. Our previous study validated the TIDAL conflict test and unmasked sex differences in the salience of anxiety- vs. pain-related stimuli that parallel clinical prevalence of anxiety; i.e., females (vs. males) exhibit increased anxiety-like behavior (Lee et al., 2022). Here, we test female and male mice with moderate SCI to reveal the salience of anxiety vs. thermal sensitivity in mice with neurotrauma. With increasing temperature in the TIDAL test, it was unclear how mice with SCI would behave: would mice leave the heated plate more quickly following neurotrauma, suggesting high salience of heat hypersensitivity; or would they persist on the dark heated plate, implying increased salience of anxiety? We find that mice with SCI more strongly prefer the dark-heated plate – even at more aversive heated temperatures. This did not appear to merely be driven by a decrease in sensitivity to heat following SCI as behavior differed from results in a thermal preference test. Overall, our results suggest that SCI amplifies the salience of an anxiety-inducing stimulus (vs. heat).

## Materials and Methods

### Animals and housing

All housing, surgery, and postoperative care were approved by The University of Texas at Austin Institutional Animal Care and Use Committee. All animals were fed standard chow and filtered tap water *ad libitum* and maintained on a 12:12 light/dark cycle. Adult (8-12 weeks old) male and female C57BL/6J mice (Jackson stock 000664) were tested during the light cycle. Mice were housed in pairs. Mice in all treatment groups were numbered randomly to ensure researchers were blind to group. At the experimental endpoint, mice were injected with an overdose of Pentobarbital (200-270 mg/kg, MWI Animal Health 011355) and tissue was collected for potential later analyses.

### Behavioral tests for anxiety-like behavior

#### Thermal Increments Dark-Light (TIDAL) Conflict Test

The TIDAL conflict apparatus is a modified thermal place preference (TPP) apparatus (Ugo Basile, Cat. No. 35250), which consists of two cylinders (20 cm diameter x 25 cm high) connected by a narrow center walkway (**Fig. 1**). For TIDAL testing, one cylinder (the “light chamber”) is kept in constant light and at a temperature of 31°C, which is an isothermic temperature for mice; in contrast, the other cylinder (“dark chamber”) is covered with a fitted opaque lid and a flexible opaque outside cover to maintain darkness inside the cylinder and the temperature is manipulated from 31 to 44°C (**Fig. 1A,C**). Light cylinder illumination levels were 550 lux, and dark cylinder illumination levels were 8 lux. In addition, the center walkway was covered with a clear plastic film “roof” to limit mouse interest in escaping through the open space. Mice are not acclimated to the apparatus prior to testing. To optimally detect the salience of anxiety vs. thermal avoidance we defined the following parameters: Mice are initially allowed to explore the apparatus for 5 minutes with both plates at 31°C (exploratory phase; initial light-dark test); next, an additional five minutes is spent with both plates at 31°C; then, the temperature on the dark plate is raised to 39°C and increased by 1°C every five minutes to a maximum temperature of 44°C (with the light plate maintained at an isothermic 31°C) (**Fig. 1A-D**).

**Figure 1.**
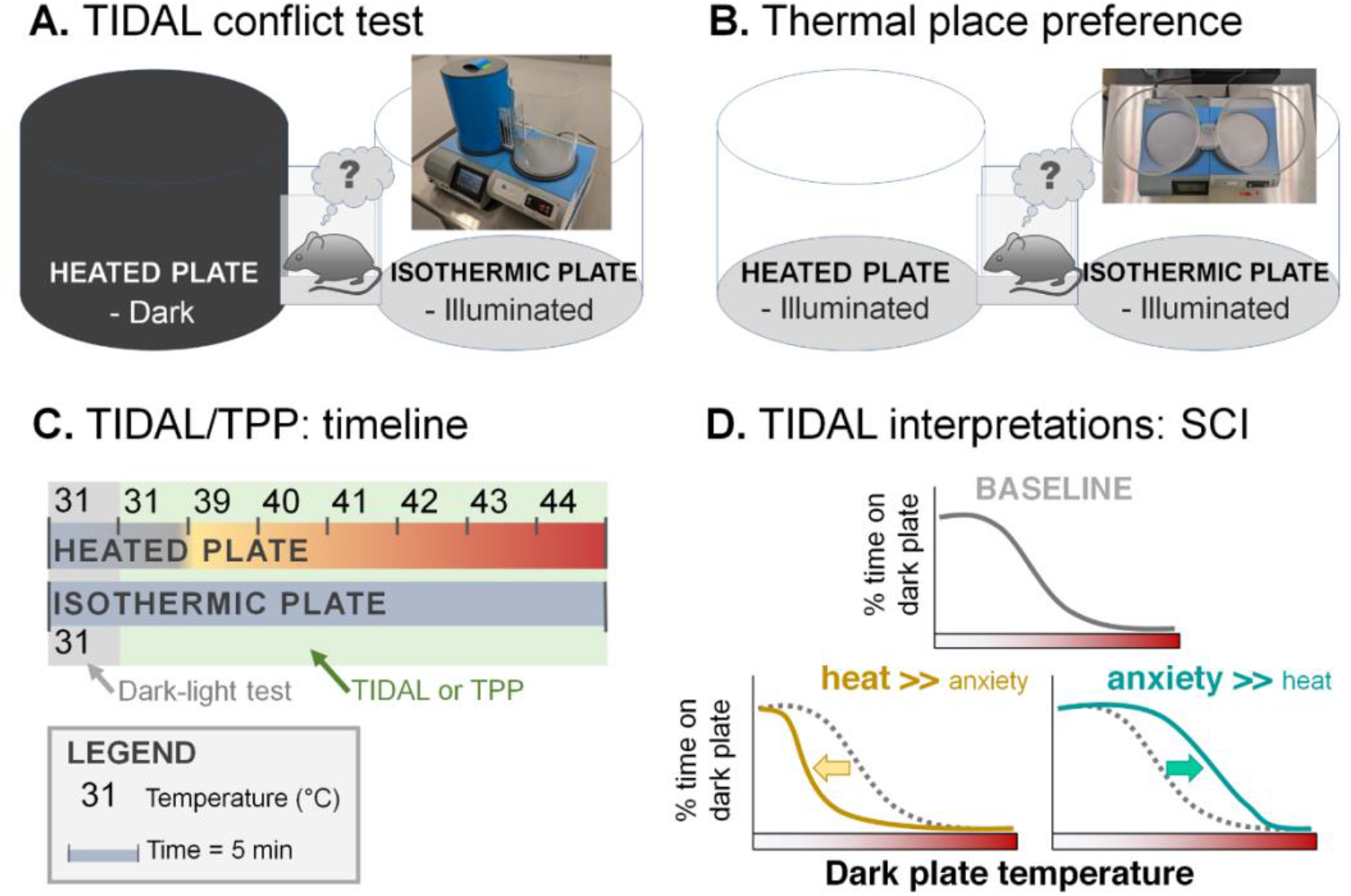
The TIDAL conflict test is a new assay for exploring the relative salience of anxiety vs. heat sensitivity. **A.** The TIDAL conflict test apparatus consists of two temperature-controlled plates linked by a center walkway. The mouse can freely move between an illuminated plate, which remains at an isothermic 31°C, and a dark plate, which starts at 31°C but incrementally increases to aversive temperatures. **B.** The thermal place preference (TPP) test is a control condition for the TIDAL test. Both chambers in TPP are illuminated; one plate increases in temperature, while the other plate remains at an isothermic 31°C. **C.** Timeline of an TIDAL/TPP test, which takes ~40 min per mouse. The first 5 min with both plates at 31°C is an acclimation period and is a control dark-light test. The TIDAL/TPP test begins with a second 31°C session, then the heated plate increases in temperature incrementally, 1°C every 5 min, from 39 to 44°C. The isothermic plate remains at 31°C throughout the test. **D.** Interpretation of potential TIDAL outcomes after SCI. At baseline (e.g., uninjured), mice are expected to decrease dark plate preference as its temperature increases. If SCI increases salience of heat (relative to dark-light), mice will leave the dark plate earlier – i.e., shifting the curve to the left. If SCI increases salience of anxiety (relative to heat), mice will prefer the dark plate to higher temperatures - i.e., shifting the curve to the right.

#### Thermal Place Preference (TPP) Assay

The TPP assay is used as a control to isolate thermal sensitivity from the anxiety-like portion of the TIDAL conflict assay. The TPP setup is the same as the TIDAL setup (two cylinders connected by a center walkway), except that both the heated side and the side maintained at 31°C are exposed to room lighting (**Fig. 1B**) – i.e., the heated side is illuminated, not dark as in the TIDAL conflict assay. Next, the same incremental temperature increases are initiated.

#### TIDAL and TPP – testing, automated video recording, and analysis

Mice tested on TPP and TIDAL assays were interspersed throughout the day (i.e., during the light phase – Zeitgeber time 1-11). Unless otherwise noted, distinct mice were used for these tests to avoid effects of learning observed in repeated testing. The percent time spent in the dark cylinder (dark cylinder time/(dark + illuminated time) – center walkway time excluded), distance traveled, and dark crossings were automatically recorded and scored using an overhead video camera and EthoVision software. Time in the center walkway was excluded from analyses in the main manuscript for two reasons: (1) the surroundings in the center zone differed from the test chambers; and (2) analyzing behavior in the identically-shaped illuminated chamber vs. dark-heating chamber enabled a two-chamber preference comparison with equal preference clearly defined at 50% time in each chamber. Data including the center zone is presented in the corresponding Supplementary figures, and show that including the center zone in analysis has little effect on the percent dark plate preference differences between groups. The arena was cleaned with 70% ethanol between trials.

### Surgery and locomotor testing

#### Surgeries – laminectomy and spinal cord injury

For all experiments, mouse surgeries were interspersed throughout the day (during the light cycle). Male and female mice were anesthetized with isoflurane inhalation anesthesia (1.5%; MWI Animal Health 502017) and treated with prophylactic buprenorphine (0.075 mg/kg; MWI Animal Health 060969) immediately prior to surgery. A dorsal T9 laminectomy was performed. The periosteum, but not the dura, was removed for all surgeries (this is the end of the surgery for sham mice). SCI mice were then subjected to a moderate contusion SCI (65 kDyn, 0 s dwell) at thoracic level 9 (T9) using the Infinite Horizon impactor (Precision Systems and Instrumentation) (Gaudet et al., 2016). SCI mice from TPP and TIDAL groups had similar injury force and displacement, respectively: female-TPP: 67.3 ± 0.9 kDyn, 524 ± 17 μm; female-TIDAL: 68 ± 2 kDyn, 524 ± 13 μm; male-TPP: 68 ± 1 kDyn, 521 ± 30 μm; male-TIDAL: 68 ± 1 kDyn, 516 ± 36 μm. Mice were monitored daily for infection or signs of abnormal recovery. To limit confounding effects on sensitivity-related behaviors, post-surgery analgesics were withheld. Post-operative mouse care included manual voiding of bladders twice daily, and hydration support via daily subcutaneous injection of Ringer’s solution (2, 2, 1, 1, 1 mL on the first 5 days post-operative (dpo), respectively).

#### Locomotor testing (Basso Mouse Scale)

Mouse locomotion was assessed before surgery and at 1, 4, 7, 10, and 14 dpo using the Basso Mouse Scale (BMS) (Basso et al., 2006). Movement of mouse hindlimbs and walking were assessed for four minutes in an open field by two condition-blind observers. Scores on the scale range from 9 (healthy mouse with coordinated, parallel steps and trunk stability) to 0 (no movement of ankle joints).

### Experiments and mouse numbers

Mice were 6-8 weeks old at time of testing. Groups of mice included female-uninjured-TPP (n=13), female-uninjured-TIDAL (n=17), female-SCI-TPP (n=9), female-SCI-TIDAL (n=17), male-uninjured-TPP (n=7), male-uninjured-TIDAL (n=24), male-SCI-TPP (n=9), and male-SCI-TIDAL (n=19). Several mice died after surgery and prior to TPP/TIDAL testing, including 5 that died during or immediately after surgery; 5 that were euthanized due to poor recovery after surgery; and 2 that were found dead in their cage. No differences were observed between naïve and sham mice, so these groups were combined as “uninjured” (6 naïve females and 6 naïve females in the TIDAL groups; all the rest received sham surgery). A subset of 7 dpo TIDAL mice were re-tested on TIDAL at 21 dpo: female-uninjured (n=12), female-SCI (n=8), male-uninjured (n=12), and male-SCI (n=9).

### Statistics

Mouse TIDAL behavior (dark plate preference, distance traveled) was analyzed using one-, two-, or three-way ANOVA (repeated measures, when appropriate), followed by Bonferroni *post-hoc* tests. In experiments with two groups, a Student’s *t-*test (or nonparametric Mann–Whitney *U* test) was performed. Prism 9 (GraphPad) was used for visualizing data and Sigmaplot 14 (SPSS) was used for statistical analyses.

## Results

### In the TIDAL conflict test, spinal cord injury increases anxiety-like behavior

To assess the salience of anxiety vs. thermal sensitivity in mice with neurotrauma, we tested male and female mice with SCI using the TIDAL conflict test or control TPP (see potential effects of SCI on TIDAL behavior in **Fig. 1D**). Mice received moderate SCI or were control uninjured mice. Mice with SCI from both groups had similar locomotor recovery from 1 to 14 dpo (Basso Mouse Scale (BMS) scores, **Fig. 2**), suggesting that differences in plate preferences between SCI mice in the TPP and TIDAL tests were solely due to the illuminated vs. dark heated plate (not due to group differences in locomotor recovery).

**Figure 2.**
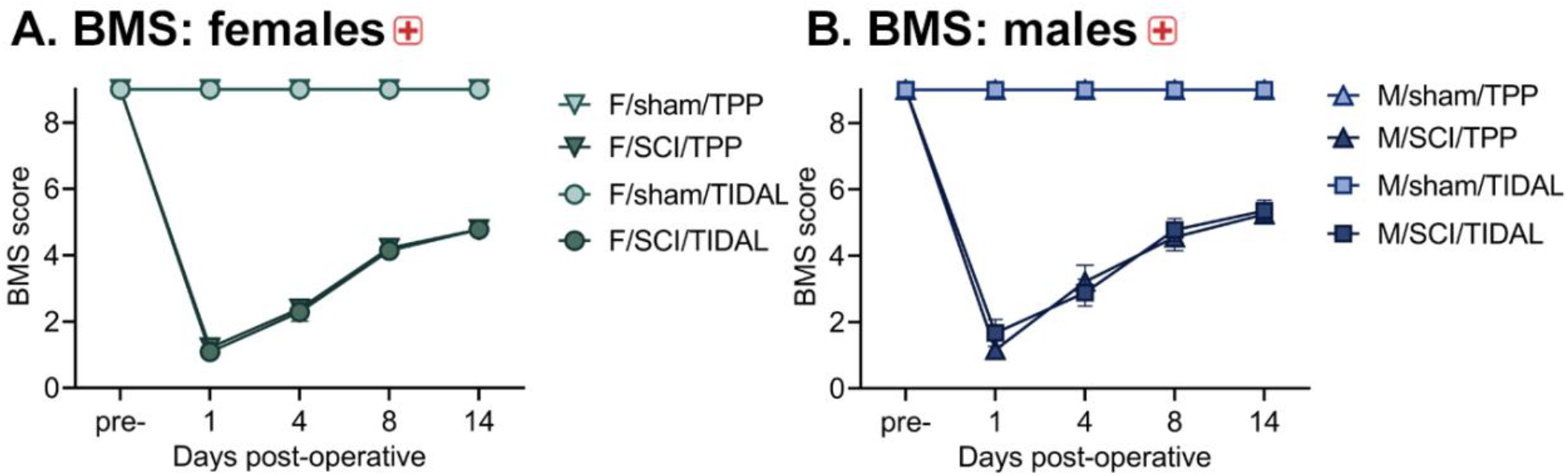
BMS locomotor recovery after SCI or sham surgery. Moderate T9 SCI caused expected locomotor deficits in female and male mice that recovered over time, as assessed in an open field using the BMS scale (**A**: females, **B**: males). Mice that received sham surgery maintained intact locomotor function. There were no significant differences in BMS scores between female and male SCI mice. Red cross symbol indicates significant main effect of surgery.

Separate cohorts of uninjured and SCI mice were tested at 7 dpo on the TIDAL conflict test or TPP test. The TPP test is important to include, because thermal preference may shift due to SCI-elicited differences in neuropathic pain-related behavior or body temperature regulation (Gaudet et al., 2017; Gaudet et al., 2018; Gaudet et al., 2021; McFarlane et al., 2020; Price and Trbovich, 2018).

In the dark-light test, mice showed robust dark plate preference vs. the illuminated plate (all groups averaged more than 68% dark plate preference), whereas mice tested with both plates illuminated showed no preference for the equivalent-but-illuminated plate (three-way ANOVA, significant interaction between test x surgery; *F*_1,106_=4.07, *p* < 0.05) (**Fig. 3A**). Mice with SCI had higher preference for the isothermic dark plate vs. uninjured mice (dark plate preference with sexes grouped: uninj., 72 ± 2%; SCI, 77 ± 2%). There were no surgery group differences with both plates illuminated. There were no significant sex differences. Thus, the dark-light test did not reveal significant differences in preference for the dark plate due to sex and exposed modestly increased anxiety-like behavior caused by SCI, suggesting that the TIDAL conflict test could better uncover surgery- and sex-related differences in anxiety-like behavior.

**Figure 3.**
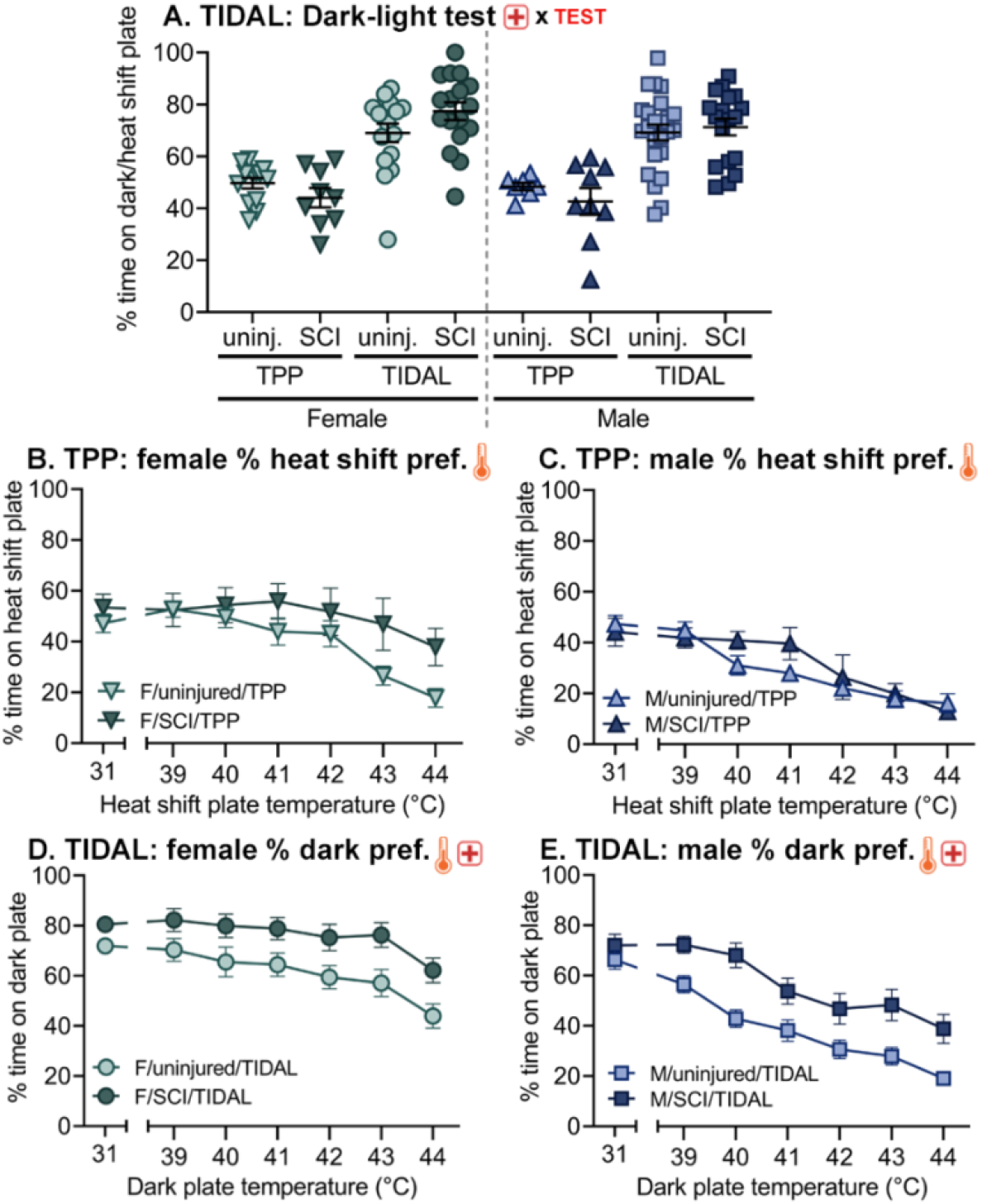
Behavior in the Thermal Increments Dark-Light (TIDAL) conflict test, compared to the control thermal place preference (TPP) test with both sides illuminated. SCI causes mice in the TIDAL conflict test to spend more time on the dark, heated plate, suggestive of anxiety-like behavior at 7 d post-SCI. **A.** In the dark-light test (TIDAL) or control light/light test (TPP) with both plates at 31°C, mice spent longer on the heated plate if it was maintained in darkness. In TIDAL, SCI mice spent more time on the heated, dark plate compared to sham mice. In contrast, in TPP conditions, mice with SCI or sham surgery showed no significant differences. **B,C.** In TPP with both plates illuminated, both females (**B**) and males (**C**) decreased heated plate preference with increasing temperature. For TPP, there was no significant effect of SCI. **D,E.** In TIDAL, both female (**D**) and male (**E**) mice with SCI had increased preference for the dark plate compared to uninjured controls. * indicates *p* < 0.05 between female and male mice; thermometer, TEST, or red cross symbols alone indicate significant main effects of temperature, TPP/TIDAL, and surgery, respectively.

Next, we explored whether the TIDAL conflict test exposes anxiety-like behavior at 7 d post-SCI. Accordingly, uninjured and SCI mice experienced increasing temperatures in the TPP and TIDAL tests. In the TPP test, female and male mice gradually decreased preference for the heat shift plate as temperatures increased (two-way RM ANOVA, main effect of temperature; both females and males *p* < 0.001), and SCI had no significant effect on TPP behavior (**Fig. 3B,C**). On the TIDAL conflict test, both female and male mice with SCI showed amplified dark plate preference (vs. uninjured; two-way RM ANOVA, main effect of surgery) (females: *F*_1,192_=7.88, *p* = 0.008) (males: *F*_1,240_=13.18, *p* < 0.001) **(Fig. 3D,E)**. Further, mice with SCI showed increased percent crossings into the dark chamber in TIDAL and traveled a similar distance compared to mice with sham surgery (**Fig. S1**).

### The TIDAL conflict test exposes differences in anxiety caused by sex and by SCI

At 40°C – a TIDAL temperature that showed notable group differences – SCI increased preference for the dark/heat shift plate (three-way ANOVA, main effect of SCI, *F*_1,106_=12.66, *p* = 0.008) (**Fig. 4A**). In addition, females increased preference for the dark and/or heated plate at 40°C compared to males, and mice on TIDAL increased preference for the heat shift plate compared to mice completing TPP (both main effects). Next, the temperature at which mice spent <50% on the heated plate was assessed. SCI increased the <50% heated plate preference temperature on TIDAL, but not TPP (<50% heated plate preference: F-uninj-TPP, 40.1 ± 0.5°C; F-SCI-TPP, 40.7 ± 0.6°C; F-uninj-TIDAL, 42.6 ± 0.4°C; F-SCI-TIDAL, 44.0 ± 0.4°C; M-uninj-TPP, 39.1 ± 0.7°C; M-SCI-TPP, 39.1 ± 0.6°C; M-uninj-TIDAL: 40.3 ± 0.4°C M-SCI-TIDAL: 41.9 ± 0.4°C) (three-way ANOVA, surgery x test interaction; *F*_1,106_=4.39, *p* < 0.05) (**Fig. 4B**). In addition, females had higher <50% heated plate preference temperature (main effect).

**Figure 4.**
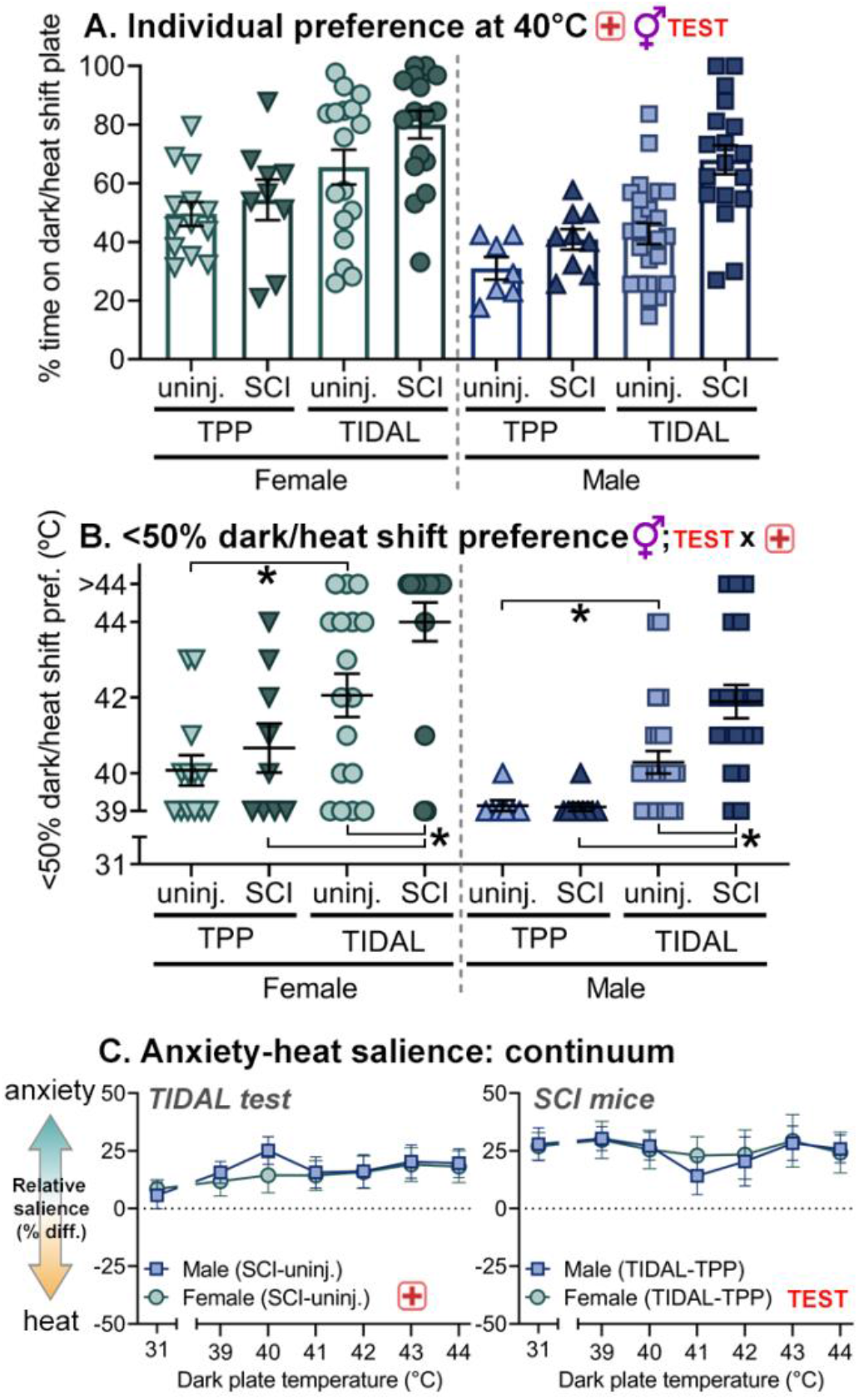
In the TIDAL conflict test, SCI increases preference for the dark, heated plate, implying increased salience of anxiety over heat sensitivity. **A.** Individual preference of mice at 40°C shows that SCI increases preference for the heated and dark plate. In addition, females (vs. males) and TIDAL (vs. TPP) had higher dark plate preferences. **B.** SCI increased the <50% dark plate preference threshold on the TIDAL test only. In addition, females had higher 50% preference temperature than males (main effect). **C.** Anxiety-heat salience continuum. Difference scores were calculated to better delineate differences between surgery and test groups. Left panel: Subtracting uninjured from SCI percent dark plate preference on TIDAL, both females and males show significantly increased salience of the anxiety-inducing stimulus (dark) vs. heat. Right panel: Subtracting TPP from TIDAL percent dark plate preference, SCI mice on TIDAL more strongly prefer the heated, dark plate compared to SCI mice completing TPP. “TEST x red cross” symbol indicates significant TPP/TIDAL x surgery interaction; gender, TEST, or red cross symbols alone indicate significant main effects of sex, TPP/TIDAL, and surgery, respectively.

To further assess the relative salience of anxiety vs. heat hypersensitivity, difference scores were calculated (**Fig. 4C**). For TIDAL, subtracting SCI minus sham scores, SCI elicited anxiety-like behavior (three-way ANOVA, main effect of surgery, *F*_1,82_=20.30, *p* < 0.0001). Averaging from 39-44°C, female-SCI mice showed 16% increased dark plate preference and male-SCI mice had 19% increased dark plate preference compared to uninjured controls. For SCI, subtracting TIDAL minus TPP scores, SCI-TIDAL mice increased dark place preference and thus anxiety-like behavior (vs. TPP controls – F-SCI-TIDAL, 39-44°C: 26% higher; M-SCI-TIDAL: 24% higher) (three-way ANOVA, main effect of test, *F*_1,49_=36.07, *p* < 0.0001). This increased dark plate preference for mice with SCI – despite rising temperatures – suggests that neurotrauma amplifies the salience of anxiety (vs. heat). Together, these data imply that SCI increases anxiety-like behavior, and that the TIDAL conflict test effectively reveals the salience of anxiety in models of neurotrauma.

### Repeating TIDAL conflict testing at 7 and 21 d post-SCI suggests interaction between learning and recovery

Next, we sought to determine whether mice with prior exposure to TIDAL show evidence of learning. Repeating the test at two distinct post-operative timepoints – 7 dpo and 21 dpo – would be informative for two reasons: (1) it would reveal whether mice learn the test and behave differently upon subsequent exposures; and (2) it would inform about anxiety-like behavioral responses at acute and near-chronic timepoints.

Mice completed the dark-light test twice; once upon first exposure to the apparatus at 7 dpo and once 14 d later after completing the entire TIDAL protocol at 21 dpo. In females completing the dark-light test, there were no significant differences in dark preference between uninjured and SCI mice at 7 or 21 dpo (*p* = 0.22) (**Fig. 5A**).

**Figure 5.**
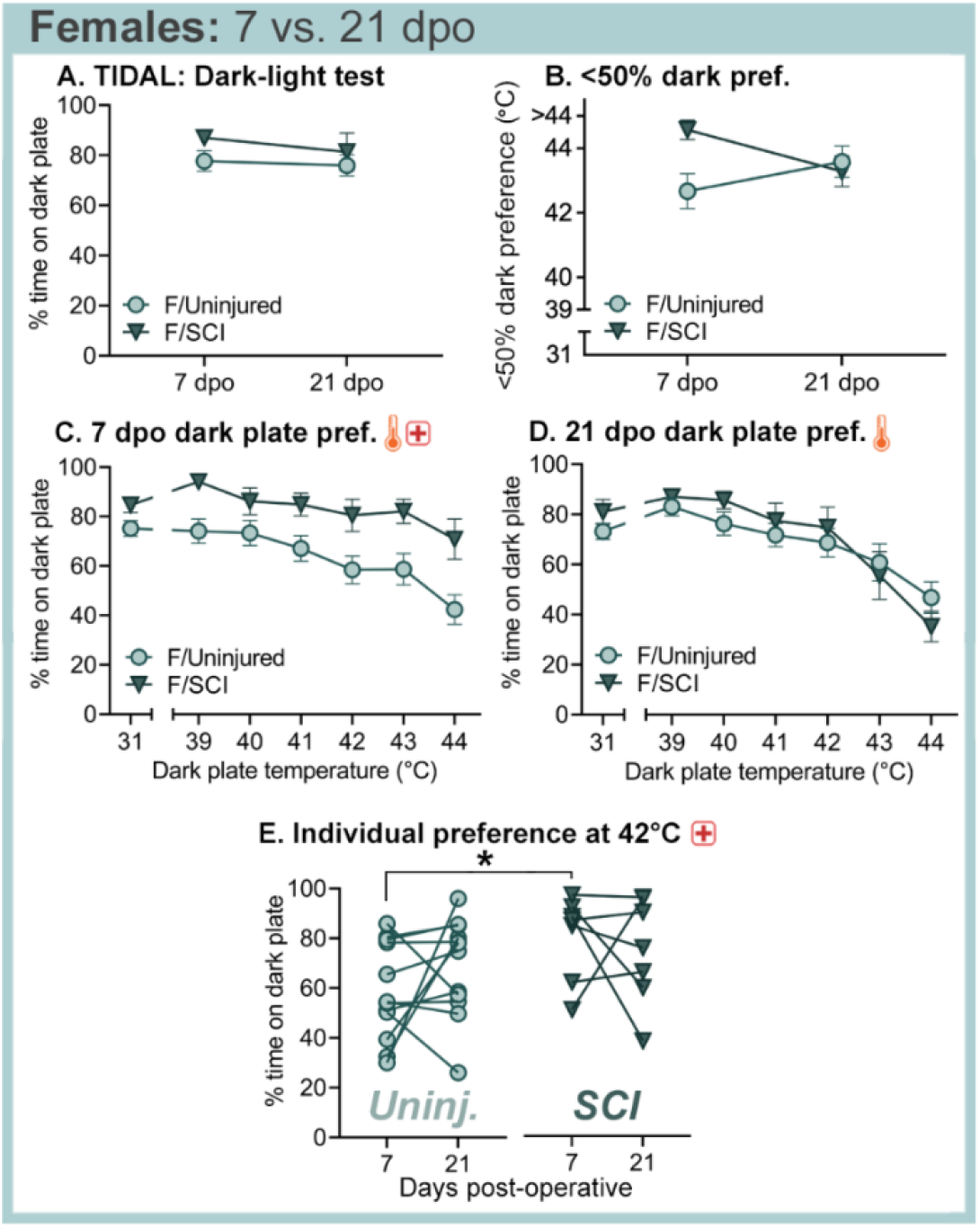
Over two sessions of TIDAL conflict tests, female mice with sham surgery shift behavior by remaining on the dark, heated side as the temperature rises. At 21 dpo, female SCI mice exhibit similar behavior as at 7 dpo, but leave more rapidly at higher temperatures. **A.** In the dark-light test with both plates at 31°C, injury had no significant effect on dark plate preferences at 7 or 21 dpo (although SCI mice tended to stay to higher temperatures at 7 dpo; surgery x dpo interaction: *p* = 0.054). **B.** Threshold at which female mice showed <50% preference for the dark plate. Injury had no significant effect on overall dark plate preference. **C, D.** Female uninjured and SCI mice were tested at 7 dpo and 21 dpo in the TIDAL conflict test with increasing temperature on the dark plate only. Female SCI (vs. uninjured) mice showed increased dark plate preference throughout the TIDAL conflict test at 7 dpo (**C**). When repeated at 21 dpo, female mice with SCI no longer show significantly increased dark plate preference compared to uninjured mice (**D**). **E-F.** Dark plate preferences of individual uninjured and SCI mice with the dark plate at 42°C. At 7 dpo, SCI females showed increased dark plate preference at 42°C relative to uninjured females (7 dpo uninjured: 56%; 7 dpo SCI: 81%). *n=12* uninjured female, *n*=8 female SCI * indicates *p* < 0.05 between SCI and uninjured mice; thermometer and red cross symbols indicate significant main effects of temperature and surgery, respectively.

At 7 dpo, female mice with SCI persisted on the heated, dark plate to higher temperatures than uninjured females, suggesting that SCI increased the salience of anxiety (vs. heat) (7 dpo SCI vs. uninjured; two-way RM ANOVA) (main effect of surgery: *F*_1,108_=5.51, *p* < 0.05) (**Fig. 5C**, **Fig. S4**). SCI mice also had increased percent crossings into the dark plate compared to sham mice (**Fig. S2**). Interestingly, at 21 dpo, there were no significant differences in dark plate preference between female uninjured and SCI mice (*p* = 0.91) (**Fig. 5D**, **Fig. S2**, **Fig. S5**). In particular, the uninjured mice more strongly preferred the dark plate at 21 dpo compared to 7 dpo, suggesting that at lower heated temperatures they learned to prefer the dark plate and exhibited increased anxiety-like behavior. At higher temperatures (43-44°C), both uninjured and SCI females showed reduced preference for the dark plate at 21 dpo vs. 7 dpo, suggesting that they anticipated the aversive nature of these higher temperatures.

At a notable heated temperature, 42°C, SCI caused increased preference for the dark plate (two-way RM ANOVA, main effect of surgery, *F*_1,37_=4.51, *p* < 0.05) (**Fig. 5E**). SCI had a particularly robust effect on 42°C dark plate preference at 7 dpo: SCI mice at 7 dpo had a stronger preference for the dark plate compared to uninjured mice (7 dpo dark plate preference: uninj., 58 ± 6%; SCI, 80 ± 6%; *p* < 0.05), whereas SCI and uninjured mice at 21 dpo had more similar 42°C dark plate preferences (21 dpo dark plate preference: uninj., 69 ± 6%; SCI, 75 ± 8%; *p* = 0.49).

Male mice with SCI and uninjured controls also completed the TIDAL conflict test at 7 and 21 dpo. In the dark-light test at 7 and 21 dpo, male mice with SCI showed no significant difference in dark-heated plate preference compared to uninjured mice (*p* = 0.28) (**Fig. 6A**). At 7 dpo, males with SCI showed a trend for increased preference for the dark plate compared to uninjured mice, although this was not significantly different (*p* = 0.14) (**Fig 6C**). At 21 dpo, uninjured male mice exhibited learning by leaving the heated, dark plate more rapidly; in contrast, male SCI mice persisted on the dark plate in a pattern that mirrored 7 dpo SCI results (21 dpo SCI vs. uninjured; two-way RM ANOVA) (significant surgery x temperature interaction: *F*_6,146_=2.47, *p* < 0.05; SCI dark preference increased at 40-42°C) (**Fig. 6D**, **Fig. S3, Fig. S5**). These results are underscored by considering a key temperature, 42°C: uninjured male mice at 42°C had reduced percent time on the dark plate at 21 dpo compared to 7 dpo (uninjured male dark plate preference: 7dpo, 33 ± 7%; 21 dpo, 17 ± 4%; *p* < 0.05) (**Fig. 6E**). Further, male mice with SCI at 42°C had increased percent time on the dark plate at 21 dpo (but not 7 dpo) compared to uninjured mice (21 dpo dark plate preference: uninj., 17 ± 4%; SCI, 44 ± 9%; *p* < 0.005). These results suggest that uninjured male mice re-exposed to TIDAL shift behavior by leaving the dark-heated plate more quickly – thereby exhibiting increased salience of the heat (vs. anxiety-related) stimulus. In contrast, male mice with SCI persist on the dark-heated plate to similar extents at both 7 and 21 dpo, suggesting that the anxiety-related aspect of the test is more salient for males with SCI compared to uninjured males.

**Figure 6.**
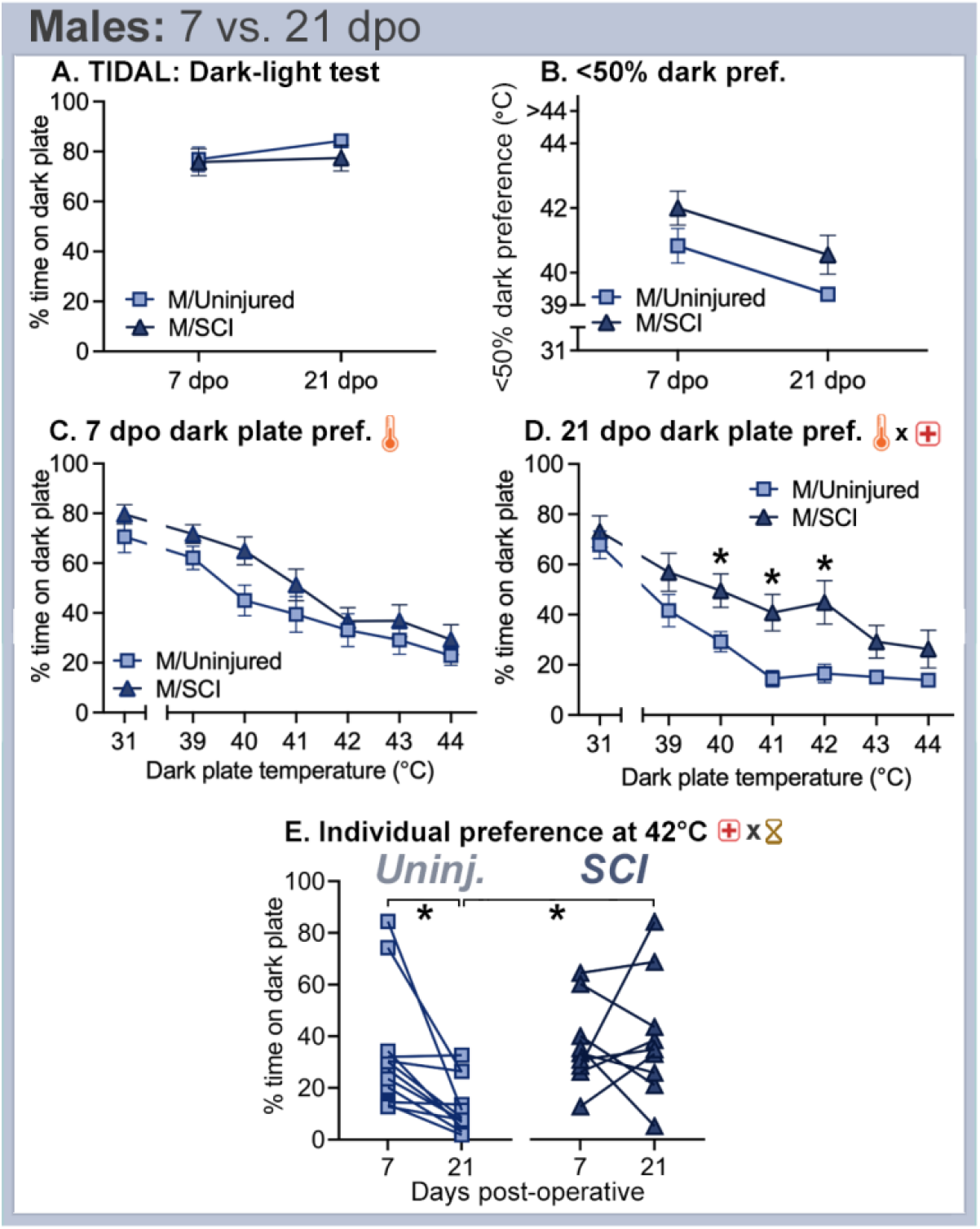
In the second session of two TIDAL conflict tests, male mice with sham surgery leave the dark-heating plate more rapidly. In contrast, male SCI mice at 21 dpo exhibit similar TIDAL behavior as at 7 dpo. **A.** In the dark-light test with both plates at 31°C, injury had no significant effect on dark plate preferences at 7 or 21 dpo. **B.** Threshold at which male mice showed <50% preference for the dark plate was decreased upon re-testing. Although SCI caused <50% preference temperatures to trend higher, injury had no significant effect on <50% dark plate preference temperature (*p* = 0.07). **C, D.** Male uninjured and SCI mice were tested at 7 dpo and 21 dpo in the TIDAL conflict test. At 7 dpo, SCI and uninjured mice had no significant difference in dark plate preference (**C**). At 21 dpi, SCI mice had increased dark plate preference between 40-42°C compared to uninjured mice (**D**). **E.** Dark plate preferences of individual uninjured and SCI mice with the dark plate at 42°C. Uninjured, but not SCI mice showed decreased dark plate preference at 21 vs. 7 dpo. At 21 dpo, SCI mice showed increased 42°C dark plate preference compared to uninjured mice. *n*=12 uninjured male, *n*=9 male SCI * indicates *p* < 0.05 between SCI and uninjured mice; thermometer and red cross symbols alone indicate significant main effects of temperature and surgery, respectively; “thermometer x red cross” symbol indicates significant temperature x surgery interaction; “red cross x hourglass” symbol indicates significant surgery x dpo interaction.

## Discussion

Here, we examined how SCI affected salience of anxiety vs. heat using a novel place preference assay, the TIDAL conflict test. In the dark-light test, uninjured and SCI mice both preferred the dark plate and there was no significant effect of SCI surgery. Uninjured mice in TIDAL left the dark plate at relatively low heated temperatures, with females (vs. males) persisting longer on the dark-heated plate. Interestingly, SCI robustly increased dark plate preference, indicative of enhanced anxiety-like symptoms. SCI increased anxiety-like behavior in TIDAL similarly for mice of both sexes. A cohort of uninjured and SCI mice were re-tested at 21 dpo; SCI mice at 21 (vs. 7) dpo showed sex-specific effects of prior testing and SCI. Whereas uninjured females re-tested on TIDAL showed increased salience of anxiety, uninjured males re-tested on TIDAL showed increased salience of temperature/heat. SCI mice of both sexes maintained similar behavior upon re-testing. Shifted anxiety-like symptoms upon re-testing likely related to a combination of learning (from prior TIDAL exposure) and sex-specific salience of anxiety vs. heat. It is notable that SCI exacerbated the salience of anxiety despite the presence of an aversive heat stimulus. Overall, our data suggest that SCI in female and male mice drives strong anxiety-like behaviors; thus, anxiety-like behavior and its related neurocircuitry could represent an understudied, therapeutically relevant target for post-SCI interventions.

### Uncovering the relative salience of anxiety- vs. pain-related stimuli after SCI

Tissue damage and functional impairment influences the perception of threat in mice. When exposed to stressful stimulation such as predator scent, injured mice avoid dangerous environments more than non-injured mice (Lister et al., 2020); in parallel, rodents with injury (here, SCI) exhibit pain-related behaviors in response to heat stimuli (Brown et al., 2021; Detloff et al., 2013; Gaudet et al., 2017; Gaudet et al., 2021; McFarlane et al., 2020). Thus, clarifying the relative salience of anxiety- vs. pain-related stimuli after SCI could aid in prioritizing research priorities and identifying neurologic mechanisms that drive comorbidities. In the light-dark test (both plates at 31°C), all groups exhibited baseline preference for the dark plate, and SCI mice had higher dark plate preference compared to uninjured mice. In the TIDAL conflict test, SCI increased dark plate preference in mice of both sexes. This was not simply due to a preference for warmth, as SCI mice on the TPP test had no (or only slight) preference for the heated side vs. sham-TPP mice, and SCI-TIDAL (vs. SCI-TPP) mice had more robust preference for the heated plate. We initially designed TIDAL/TPP as a tool to test whether mice had symptoms of anxiety and/or pain, yet our results suggest that this novel conflict test is optimized to detect anxiety-related behaviors. Indeed, the temperatures used did not expose the extent of heat-related neuropathic pain symptoms: whereas the Hargreaves test confirms that rodents with SCI display heat hyperalgesia, our TPP study (with both chambers illuminated) highlights that mice with SCI remained on the heated plate to the same or higher temperatures vs. uninjured mice.

At 21 d after SCI, mice still exhibited anxiety-like behaviors; shifting TIDAL behavior compared to 7 dpo likely reflects a combination of learning and improved neurologic recovery with time post-SCI. As mentioned above, pain researchers aim to develop preclinical tests that better incorporate the affective component of the pain experience; this study adds insight regarding the benefits and challenges of creating more complex – but more ethologically relevant – tests for thermal hypersensitivity vs. anxiety. Overall, our TIDAL conflict test data suggest that SCI in female and male mice increases anxiety-like behavior.

### SCI amplifies anxiety-like behaviors

Our work parallels previous research showing that SCI exacerbates anxiety-like behaviors in rodents and in humans. When tested on the TIDAL conflict test, mice with SCI exhibited increased anxiety-like behavior. Similarly, previous studies in rats and mice exposed SCI-elicited anxiety symptoms using single-parameter tests, such as open field, elevated plus maze, and burying behaviors (Fukutoku et al., 2020; Maldonado-Bouchard et al., 2016). Anxiety is more prevalent after SCI in humans. Given the high prevalence of anxiety disorders in individuals with SCI, it is essential to understand the underlying mechanisms responsible for decreased psychological well-being after SCI. SCI robustly increases pro-inflammatory cytokine expression and immune system activation in the central nervous system, from acute-to-chronic times post-injury (Gaudet and Fonken, 2018; Maldonado-Bouchard et al., 2016; Yip and Malaspina, 2012). In humans, elevated levels of pro-inflammatory cytokines in the nervous system correlate with anxiety (Leff Gelman et al., 2019; Miller et al., 2013). Additionally, inflammatory responses are differentially regulated in women compared to men (Takahashi and Iwasaki, 2021). Gender is a key factor in SCI outcomes; men are more likely to experience SCI, whereas women are more vulnerable to mood disorders and neuropathic pain following SCI (Wilson et al., 2018). Overall, future research must address post-SCI psychological challenges and the effect of sex; our understanding of SCI-exacerbated anxiety will be advanced through the use of effective assays for anxiety-like behavior such as conflict tests.

### Developing more translatable models and tests to explore pain-related behaviors

Ongoing research seeks to develop tests that better model chronic pain, often by incorporating affective pain-relevant behaviors. Some affective pain-relevant tests assess shifts in spontaneous rodent behavior. For example, the mouse grimace scale uses facial expression-based machine learning techniques to define rodent facial expressions in response to pain and pain-modifying agents (Heinsinger et al., 2020; Langford et al., 2010; Matsumiya et al., 2012). Chronic pain conditions can also be studied by analyzing rodent naturalistic cage behavior. Cage lid hanging behavior is a quantifiable measure of mouse pain state: mice exhibiting pain-like symptoms exhibit decreased hanging behavior, which was modified by intensity of noxious stimulation (Zhang et al., 2021). In addition, burrowing and nest building behavior are used to assess general rodent well-being, and both behaviors are impaired by stress or noxious stimulation (Deacon, 2006; Jirkof, 2014; Jirkof et al., 2010). Finally, conflict tests can be used to address affective aspects of pain-like symptoms in rodents. Conflict tests produce differing motivational states through the introduction of approach-avoidance situations; pain-like behavior can be assessed in rodents using conflict tests such as the mechanical-conflict avoidance paradigm (Chhaya et al., 2019; Gaffney et al., 2022) and conditioned place preference (Yang et al., 2014). Here, we developed the TIDAL conflict test in part to unmask SCI-elicited neuropathic pain-like states, but instead found that TIDAL is better optimized to reveal anxiety-like behaviors due to its slow increase through a finely-tuned, slightly aversive heat temperature range. Although the TIDAL conflict test does not robustly test pain-like symptoms, our data suggest that placing two ethologically relevant stimuli in conflict can help unveil significant differences in behavior across groups; future studies could leverage this information to develop new pain-relevant conflict assays and/or combine these with measures of spontaneous pain-like behaviors.

### Future directions and conclusions

In this study, our newly developed thermal increments dark-light (TIDAL) conflict assay reveals that SCI increases anxiety-like behavior. In the future, this test could help identify brain circuits and neuroinflammatory foci that underlie SCI-elicited shifts in anxiety- vs. pain-related symptoms, and to help dissect overlapping vs. independent regions involved in anxiety vs. pain. In addition, the TIDAL conflict test could be used after SCI (or for other conditions) to test new anxiolytic or analgesic drugs, to establish in a preclinical test how promising drugs affect place preference and place avoidance behavior. Further, future iterations of the TIDAL conflict test could incorporate cold (rather than hot) temperatures, or could use fewer, key temperatures to expedite testing for each mouse and to improve feasibility for larger cohorts of mice. Finally, the TIDAL conflict test exposes increased anxiety-like behavior in female mice that recapitulates sex differences in humans; future studies could use this test to examine sexual dimorphism in anxiety and pain.

In conclusion, we assessed anxiety- vs. pain-like behavior in mice with SCI using the heat-light TIDAL conflict test. Our data illuminate that SCI robustly boosts anxiety-like behavior, even as temperatures rise to aversive temperatures. Female (vs. male) mice remained on the dark-heated plate to higher temperatures, yet SCI had similar anxiety-amplifying effects in mice of both sexes. Adding an anxiety-related stimulus (dark vs. light) in conflict with an incrementally increasing heat-related stimulus (heated vs. isothermic) helped decipher the relative salience between these two related conditions. Remarkably, despite the fact that SCI elicits heat hypersensitivity as shown in previous studies, mice with SCI consistently remained on the heated-dark plate to higher temperatures than uninjured mice. The TIDAL conflict test could be useful after SCI, traumatic brain injury, and peripheral nerve injury for dissecting the relative salience of anxiety- vs. pain-related stimuli and for exploring therapeutic strategies. Overall, our data highlight that SCI increases the salience of anxiety (vs. heat), and suggest that anxiety-related pathways should be studied after SCI with an aim to ameliorate anxiety disorders and commonly co-occurring neuropathic pain.

## Funding and Disclosure

### Conflict of interest

The authors declare no competing financial interests.

## Acknowledgements

We thank the Animal Resources Center (ARC) husbandry staff at the Health Discovery Building for excellent animal care. Partial support was provided by University of Texas at Austin start-up funds (ADG), the Wings for Life Foundation (ADG) and Mission Connect, a program of the TIRR Foundation (ADG).

## Notes

### Competing Interest Statement

The authors have declared no competing interest.

## References

Anderson, K.D., 2004. Targeting recovery: priorities of the spinal cord-injured population. Journal of Neurotrauma 21, 1371–1383.

Askari, M.S., Andrade, L.H., Filho, A.C., Silveira, C.M., Siu, E., Wang, Y.-P., Viana, M.C., Martins, S.S., 2017. Dual burden of chronic physical diseases and anxiety/mood disorders among São Paulo Megacity Mental Health Survey Sample, Brazil. J Affect Disord 220, 1–7.

Bekhbat, M., Neigh, G.N., 2018. Sex differences in the neuro-immune consequences of stress: Focus on depression and anxiety. Brain, Behavior, and Immunity 67, 1–12.

Brown, E.V., Falnikar, A., Heinsinger, N., Cheng, L., Andrews, C.E., DeMarco, M., Lepore, A.C., 2021. Cervical spinal cord injury-induced neuropathic pain in male mice is associated with a persistent pro-inflammatory macrophage/microglial response in the superficial dorsal horn. Experimental Neurology 343, 113757.

Burma, N.E., Leduc-Pessah, H., Fan, C.Y., Trang, T., 2017. Animal models of chronic pain: Advances and challenges for clinical translation. Journal of Neuroscience Research 95, 1242–1256.

Chhaya, S.J., Quiros-Molina, D., Tamashiro-Orrego, A.D., Houlé, J.D., Detloff, M.R., 2019. Exercise-Induced Changes to the Macrophage Response in the Dorsal Root Ganglia Prevent Neuropathic Pain after Spinal Cord Injury. Journal of Neurotrauma 36, 877–890.

Collinger, J.L., Boninger, M.L., Bruns, T.M., Curley, K., Wang, W., Weber, D.J., 2013. Functional priorities, assistive technology, and brain-computer interfaces after spinal cord injury. Journal of Rehabilitation Research and Development 50, 145–160.

Dahan, A., van Velzen, M., Niesters, M., 2014. Comorbidities and the complexities of chronic pain. Anesthesiology 121, 675–677.

Deacon, R.M., 2006. Assessing nest building in mice. Nature Protocols 1, 1117–1119.

Detloff, M.R., Wade, R.E., Jr., Houle, J.D., 2013. Chronic at- and below-level pain after moderate unilateral cervical spinal cord contusion in rats. Journal of Neurotrauma 30, 884–890.

Fukutoku, T., Kumagai, G., Fujita, T., Sasaki, A., Wada, K., Liu, X., Tanaka, T., Kudo, H., Asari, T., Nikaido, Y., Ueno, S., Ishibashi, Y., 2020. Sex-Related Differences in Anxiety and Functional Recovery after Spinal Cord Injury in Mice. Journal of Neurotrauma 37, 2235–2243.

Gaffney, C.M., Muwanga, G., Shen, H., Tawfik, V.L., Shepherd, A.J., 2022. Mechanical Conflict-Avoidance Assay to Measure Pain Behavior in Mice. Journal of Visualized Experiments: JoVE.

Gaudet, A.D., Ayala, M.T., Schleicher, W.E., Smith, E.J., Bateman, E.M., Maier, S.F., Watkins, L.R., 2017. Exploring acute-to-chronic neuropathic pain in rats after contusion spinal cord injury. Experimental Neurology 295, 46–54.

Gaudet, A.D., Fonken, L.K., 2018. Glial Cells Shape Pathology and Repair After Spinal Cord Injury. Neurotherapeutics 15, 554–577.

Gaudet, A.D., Fonken, L.K., Ayala, M.T., Bateman, E.M., Schleicher, W.E., Smith, E.J., D’Angelo, H.M., Maier, S.F., Watkins, L.R., 2018. Spinal Cord Injury in Rats Disrupts the Circadian System. eNeuro 5.

Gaudet, A.D., Fonken, L.K., Ayala, M.T., Maier, S.F., Watkins, L.R., 2021. Aging and miR-155 in mice influence survival and neuropathic pain after spinal cord injury. Brain, Behavior, and Immunity 97, 6.

Gaudet, A.D., Mandrekar-Colucci, S., Hall, J.C., Sweet, D.R., Schmitt, P.J., Xu, X., Guan, Z., Mo, X., Guerau-de-Arellano, M., Popovich, P.G., 2016. miR-155 Deletion in Mice Overcomes Neuron-Intrinsic and Neuron-Extrinsic Barriers to Spinal Cord Repair. The Journal of Neuroscience 36, 8516–8532.

Heinsinger, N.M., Spagnuolo, G., Allahyari, R.V., Galer, S., Fox, T., Jaffe, D.A., Thomas, S.J., Iacovitti, L., Lepore, A.C., 2020. Facial grimace testing as an assay of neuropathic pain-related behavior in a mouse model of cervical spinal cord injury. Experimental Neurology 334, 113468.

Jirkof, P., 2014. Burrowing and nest building behavior as indicators of well-being in mice. Journal of Neuroscience Methods 234, 139–146.

Jirkof, P., Cesarovic, N., Rettich, A., Nicholls, F., Seifert, B., Arras, M., 2010. Burrowing behavior as an indicator of post-laparotomy pain in mice. Frontiers in Behavioral Neuroscience 4, 165.

Kramer, J.L., Minhas, N.K., Jutzeler, C.R., Erskine, E.L., Liu, L.J., Ramer, M.S., 2017. Neuropathic pain following traumatic spinal cord injury: Models, measurement, and mechanisms. Journal of Neuroscience Research 95, 1295–1306.

Langford, D.J., Bailey, A.L., Chanda, M.L., Clarke, S.E., Drummond, T.E., Echols, S., Glick, S., Ingrao, J., Klassen-Ross, T., Lacroix-Fralish, M.L., Matsumiya, L., Sorge, R.E., Sotocinal, S.G., Tabaka, J.M., Wong, D., van den Maagdenberg, A.M., Ferrari, M.D., Craig, K.D., Mogil, J.S., 2010. Coding of facial expressions of pain in the laboratory mouse. Nature Methods 7, 447–449.

Le, J., Dorstyn, D., 2016. Anxiety prevalence following spinal cord injury: a meta-analysis. Spinal Cord 54, 570–578.

Lee, S.E., Greenough, E.K., Oancea, P., Fonken, L.K., Gaudet, A.D., 2022. Anxiety-like behaviors in mice unmasked: Revealing sex differences in anxiety using a novel light-heat conflict test. bioRxiv, 2022.2009.2002.506410.

Leff Gelman, P., Mancilla-Herrera, I., Flores-Ramos, M., Saravia Takashima, M.F., Cruz Coronel, F.M., Cruz Fuentes, C., Pérez Molina, A., Hernández-Ruiz, J., Silva-Aguilera, F.S., Farfan-Labonne, B., Chinchilla-Ochoa, D., Garza Morales, S., Camacho-Arroyo, I., 2019. The cytokine profile of women with severe anxiety and depression during pregnancy. BMC Psychiatry 19, 104.

Lister, K.C., Bouchard, S.M., Markova, T., Aternali, A., Denecli, P., Pimentel, S.D., Majeed, M., Austin, J.S., de, C.W.A.C., Mogil, J.S., 2020. Chronic pain produces hypervigilance to predator odor in mice. Current Biology 30, R866–r867.

Lo, C., Tran, Y., Anderson, K., Craig, A., Middleton, J., 2016. Functional Priorities in Persons with Spinal Cord Injury: Using Discrete Choice Experiments To Determine Preferences. Journal of Neurotrauma 33, 1958–1968.

Maldonado-Bouchard, S., Peters, K., Woller, S.A., Madahian, B., Faghihi, U., Patel, S., Bake, S., Hook, M.A., 2016. Inflammation is increased with anxiety- and depression-like signs in a rat model of spinal cord injury. Brain, Behavior, and Immunity 51, 176–195.

Matsumiya, L.C., Sorge, R.E., Sotocinal, S.G., Tabaka, J.M., Wieskopf, J.S., Zaloum, A., King, O.D., Mogil, J.S., 2012. Using the Mouse Grimace Scale to reevaluate the efficacy of postoperative analgesics in laboratory mice. Journal of the American Association for Laboratory Animal Science: JAALAS 51, 42–49.

McFarlane, K., Otto, T.E., Bailey, W.M., Veldhorst, A.K., Donahue, R.R., Taylor, B.K., Gensel, J.C., 2020. Effect of Sex on Motor Function, Lesion Size, and Neuropathic Pain after Contusion Spinal Cord Injury in Mice. Journal of Neurotrauma 37, 1983–1990.

Miller, A.H., Haroon, E., Raison, C.L., Felger, J.C., 2013. Cytokine targets in the brain: impact on neurotransmitters and neurocircuits. Depression and Anxiety 30, 297–306.

Price, M.J., Trbovich, M., 2018. Thermoregulation following spinal cord injury. Handbook of Clinical Neurology 157, 799–820.

Siddall, P.J., Taylor, D.A., McClelland, J.M., Rutkowski, S.B., Cousins, M.J., 1999. Pain report and the relationship of pain to physical factors in the first 6 months following spinal cord injury. Pain 81, 187–197.

Takahashi, T., Iwasaki, A., 2021. Sex differences in immune responses. Science (New York, N.Y.) 371, 347–348.

Von Korff, M., Simon, G., 1996. The relationship between pain and depression. The British Journal of Psychiatry. Supplement, 101–108.

Wilson, C.S., Nassar, S.L., Ottomanelli, L., Barnett, S.D., Njoh, E., 2018. Gender differences in depression among veterans with spinal cord injury. Rehabilitation Psychology 63, 221–229.

Wohleb, E.S., Patterson, J.M., Sharma, V., Quan, N., Godbout, J.P., Sheridan, J.F., 2014. Knockdown of interleukin-1 receptor type-1 on endothelial cells attenuated stress-induced neuroinflammation and prevented anxiety-like behavior. The Journal of Neuroscience 34, 2583–2591.

Yang, Q., Wu, Z., Hadden, J.K., Odem, M.A., Zuo, Y., Crook, R.J., Frost, J.A., Walters, E.T., 2014. Persistent pain after spinal cord injury is maintained by primary afferent activity. The Journal of neuroscience: the official journal of the Society for Neuroscience 34, 10765–10769.

Yip, P.K., Malaspina, A., 2012. Spinal cord trauma and the molecular point of no return. Molecular Neurodegeneration 7, 6.

Zhang, H., Lecker, I., Collymore, C., Dokova, A., Pham, M.C., Rosen, S.F., Crawhall-Duk, H., Zain, M., Valencia, M., Filippini, H.F., Li, J., D’Souza, A.J., Cho, C., Michailidis, V., Whissell, P.D., Patel, I., Steenland, H.W., Virginia Lee, W.J., Moayedi, M., Sterley, T.L., Bains, J.S., Stratton, J.A., Matyas, J.R., Biernaskie, J., Dubins, D., Vukobradovic, I., Bezginov, A., Flenniken, A.M., Martin, L.J., Mogil, J.S., Bonin, R.P., 2021. Cage-lid hanging behavior as a translationally relevant measure of pain in mice. Pain 162, 1416–1425.

