## Supplementary Figures: for "Spinal cord injury in mice amplifies anxiety: a novel light-heat conflict test exposes increased salience of anxiety over heat"

### Slide 1
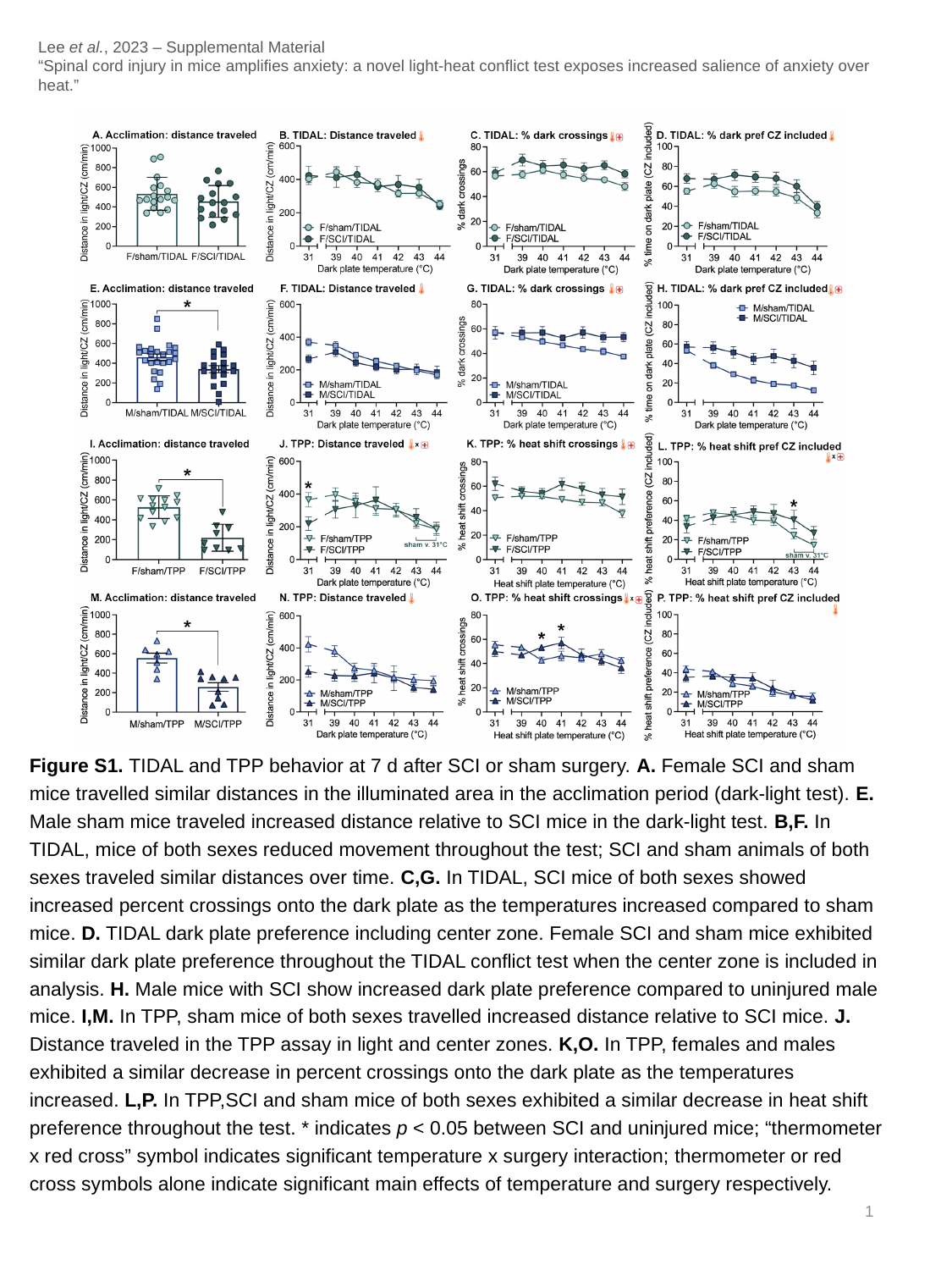

Lee et al., 2023 – Supplemental Material
“Spinal cord injury in mice amplifies anxiety: a novel light-heat conflict test exposes increased salience of anxiety over heat.”
Figure S1. TIDAL and TPP behavior at 7 d after SCI or sham surgery. A. Female SCI and sham mice travelled similar distances in the illuminated area in the acclimation period (dark-light test). E. Male sham mice traveled increased distance relative to SCI mice in the dark-light test. B,F. In TIDAL, mice of both sexes reduced movement throughout the test; SCI and sham animals of both sexes traveled similar distances over time. C,G. In TIDAL, SCI mice of both sexes showed increased percent crossings onto the dark plate as the temperatures increased compared to sham mice. D. TIDAL dark plate preference including center zone. Female SCI and sham mice exhibited similar dark plate preference throughout the TIDAL conflict test when the center zone is included in analysis. H. Male mice with SCI show increased dark plate preference compared to uninjured male mice. I,M. In TPP, sham mice of both sexes travelled increased distance relative to SCI mice. J. Distance traveled in the TPP assay in light and center zones. K,O. In TPP, females and males exhibited a similar decrease in percent crossings onto the dark plate as the temperatures increased. L,P. In TPP,SCI and sham mice of both sexes exhibited a similar decrease in heat shift preference throughout the test. * indicates p < 0.05 between SCI and uninjured mice; “thermometer x red cross” symbol indicates significant temperature x surgery interaction; thermometer or red cross symbols alone indicate significant main effects of temperature and surgery respectively.
1

### Slide 2
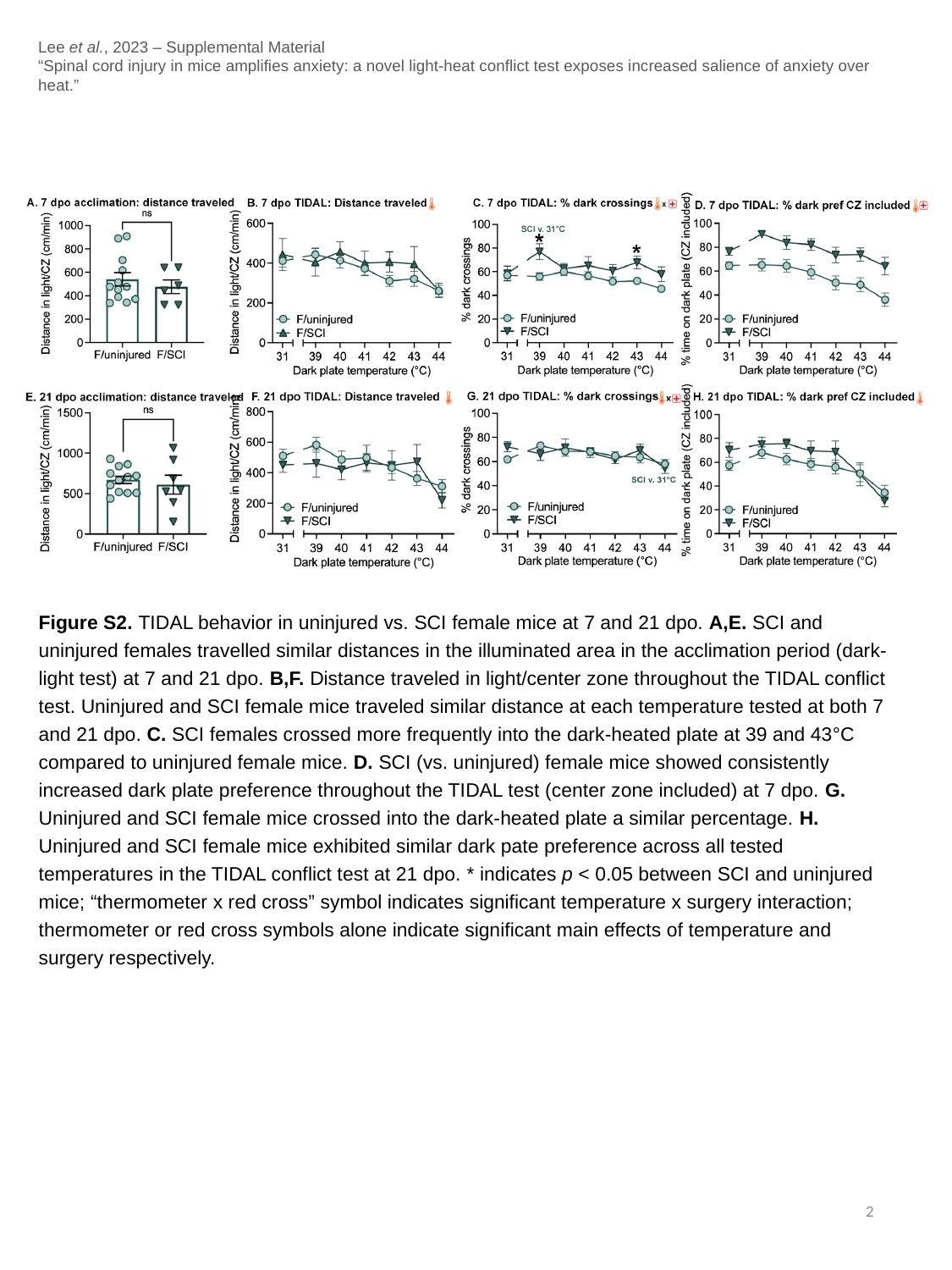

Lee et al., 2023 – Supplemental Material
“Spinal cord injury in mice amplifies anxiety: a novel light-heat conflict test exposes increased salience of anxiety over heat.”
Figure S2. TIDAL behavior in uninjured vs. SCI female mice at 7 and 21 dpo. A,E. SCI and uninjured females travelled similar distances in the illuminated area in the acclimation period (dark-light test) at 7 and 21 dpo. B,F. Distance traveled in light/center zone throughout the TIDAL conflict test. Uninjured and SCI female mice traveled similar distance at each temperature tested at both 7 and 21 dpo. C. SCI females crossed more frequently into the dark-heated plate at 39 and 43°C compared to uninjured female mice. D. SCI (vs. uninjured) female mice showed consistently increased dark plate preference throughout the TIDAL test (center zone included) at 7 dpo. G. Uninjured and SCI female mice crossed into the dark-heated plate a similar percentage. H. Uninjured and SCI female mice exhibited similar dark pate preference across all tested temperatures in the TIDAL conflict test at 21 dpo. * indicates p < 0.05 between SCI and uninjured mice; “thermometer x red cross” symbol indicates significant temperature x surgery interaction; thermometer or red cross symbols alone indicate significant main effects of temperature and surgery respectively.
2

### Slide 3
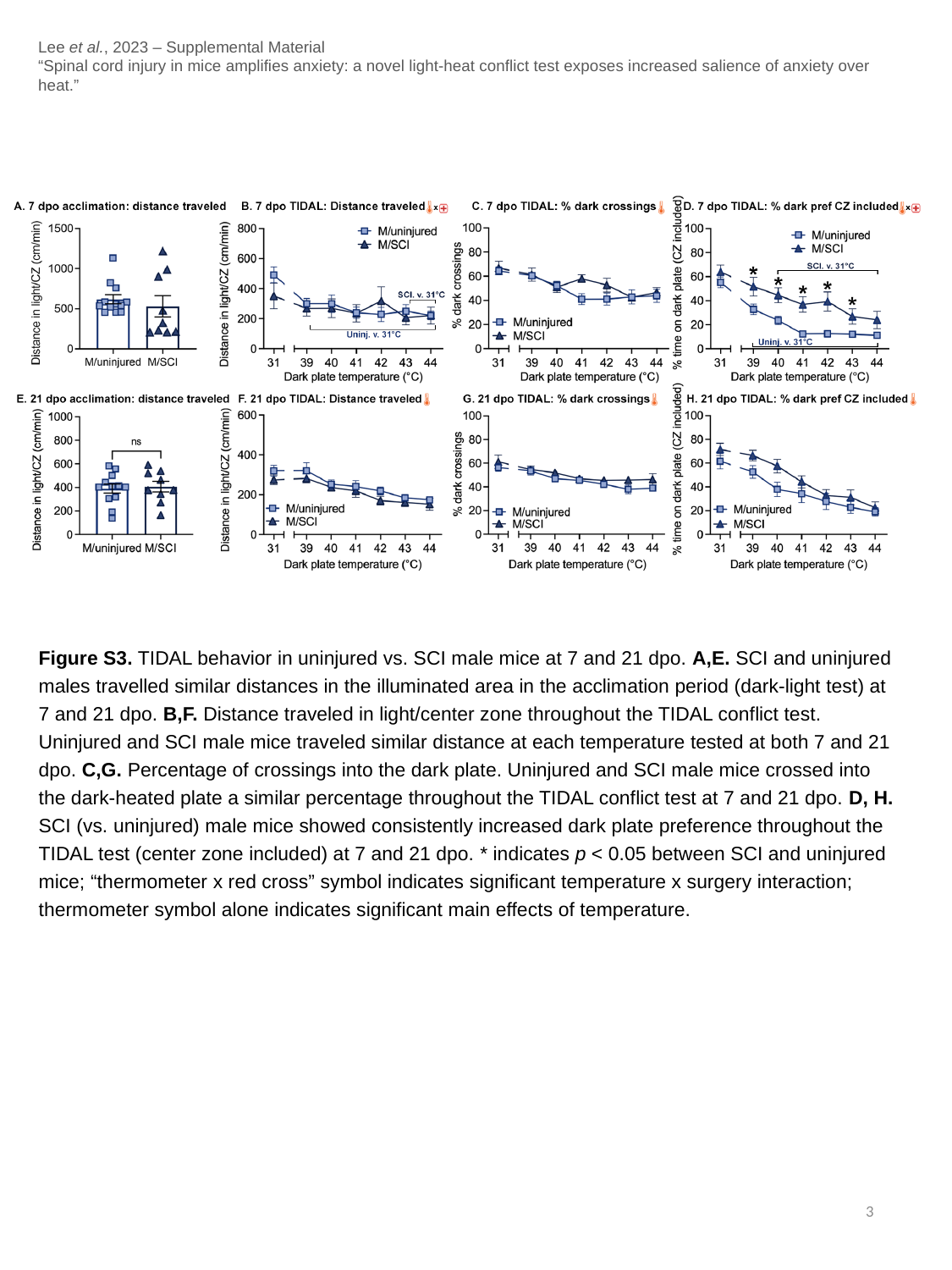

Lee et al., 2023 – Supplemental Material
“Spinal cord injury in mice amplifies anxiety: a novel light-heat conflict test exposes increased salience of anxiety over heat.”
Figure S3. TIDAL behavior in uninjured vs. SCI male mice at 7 and 21 dpo. A,E. SCI and uninjured males travelled similar distances in the illuminated area in the acclimation period (dark-light test) at 7 and 21 dpo. B,F. Distance traveled in light/center zone throughout the TIDAL conflict test. Uninjured and SCI male mice traveled similar distance at each temperature tested at both 7 and 21 dpo. C,G. Percentage of crossings into the dark plate. Uninjured and SCI male mice crossed into the dark-heated plate a similar percentage throughout the TIDAL conflict test at 7 and 21 dpo. D, H. SCI (vs. uninjured) male mice showed consistently increased dark plate preference throughout the TIDAL test (center zone included) at 7 and 21 dpo. * indicates p < 0.05 between SCI and uninjured mice; “thermometer x red cross” symbol indicates significant temperature x surgery interaction; thermometer symbol alone indicates significant main effects of temperature.
3

### Slide 4
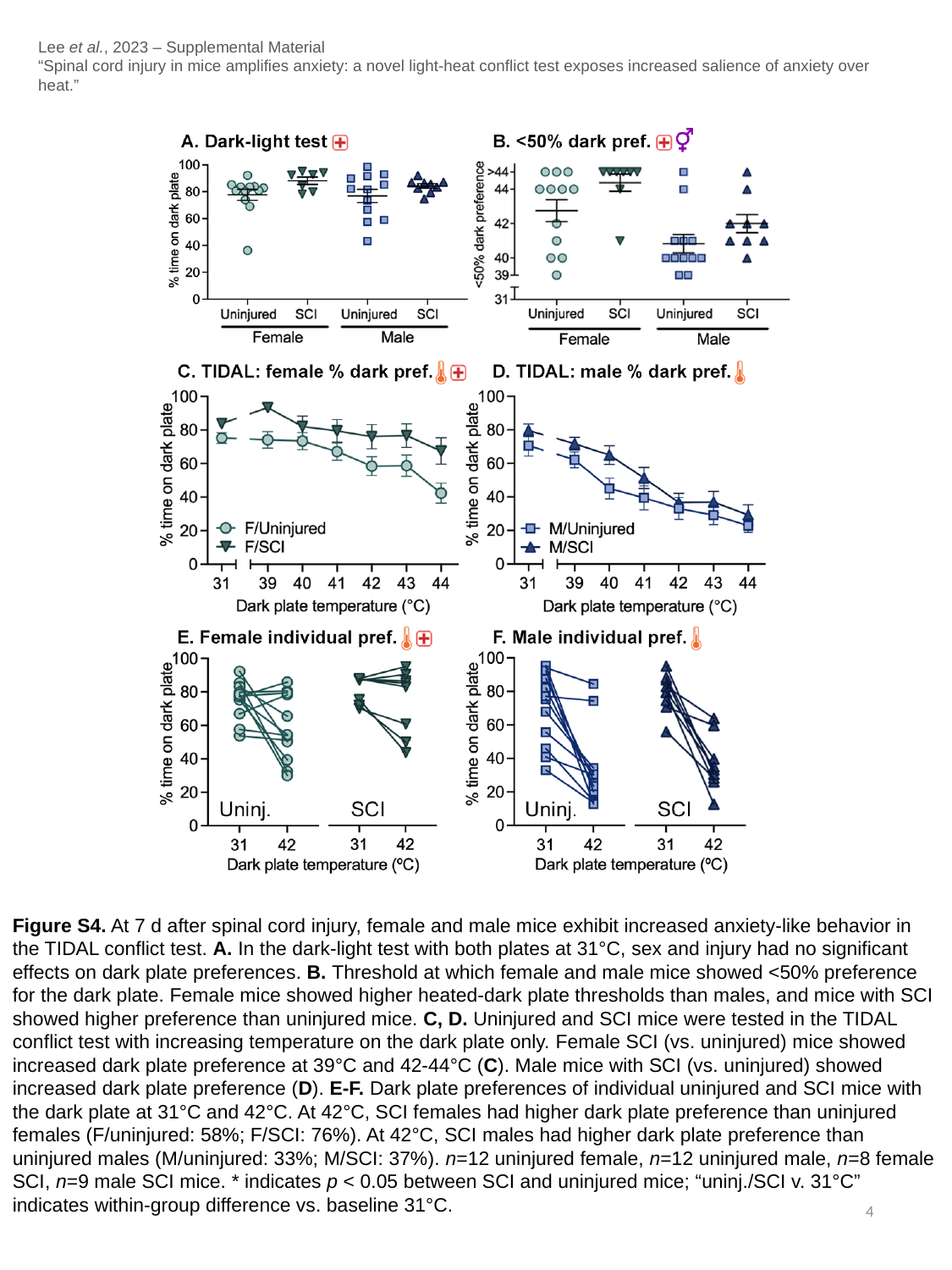

Lee et al., 2023 – Supplemental Material
“Spinal cord injury in mice amplifies anxiety: a novel light-heat conflict test exposes increased salience of anxiety over heat.”
Figure S4. At 7 d after spinal cord injury, female and male mice exhibit increased anxiety-like behavior in the TIDAL conflict test. A. In the dark-light test with both plates at 31°C, sex and injury had no significant effects on dark plate preferences. B. Threshold at which female and male mice showed <50% preference for the dark plate. Female mice showed higher heated-dark plate thresholds than males, and mice with SCI showed higher preference than uninjured mice. C, D. Uninjured and SCI mice were tested in the TIDAL conflict test with increasing temperature on the dark plate only. Female SCI (vs. uninjured) mice showed increased dark plate preference at 39°C and 42-44°C (C). Male mice with SCI (vs. uninjured) showed increased dark plate preference (D). E-F. Dark plate preferences of individual uninjured and SCI mice with the dark plate at 31°C and 42°C. At 42°C, SCI females had higher dark plate preference than uninjured females (F/uninjured: 58%; F/SCI: 76%). At 42°C, SCI males had higher dark plate preference than uninjured males (M/uninjured: 33%; M/SCI: 37%). n=12 uninjured female, n=12 uninjured male, n=8 female SCI, n=9 male SCI mice. * indicates p < 0.05 between SCI and uninjured mice; “uninj./SCI v. 31°C” indicates within-group difference vs. baseline 31°C.
4

### Slide 5
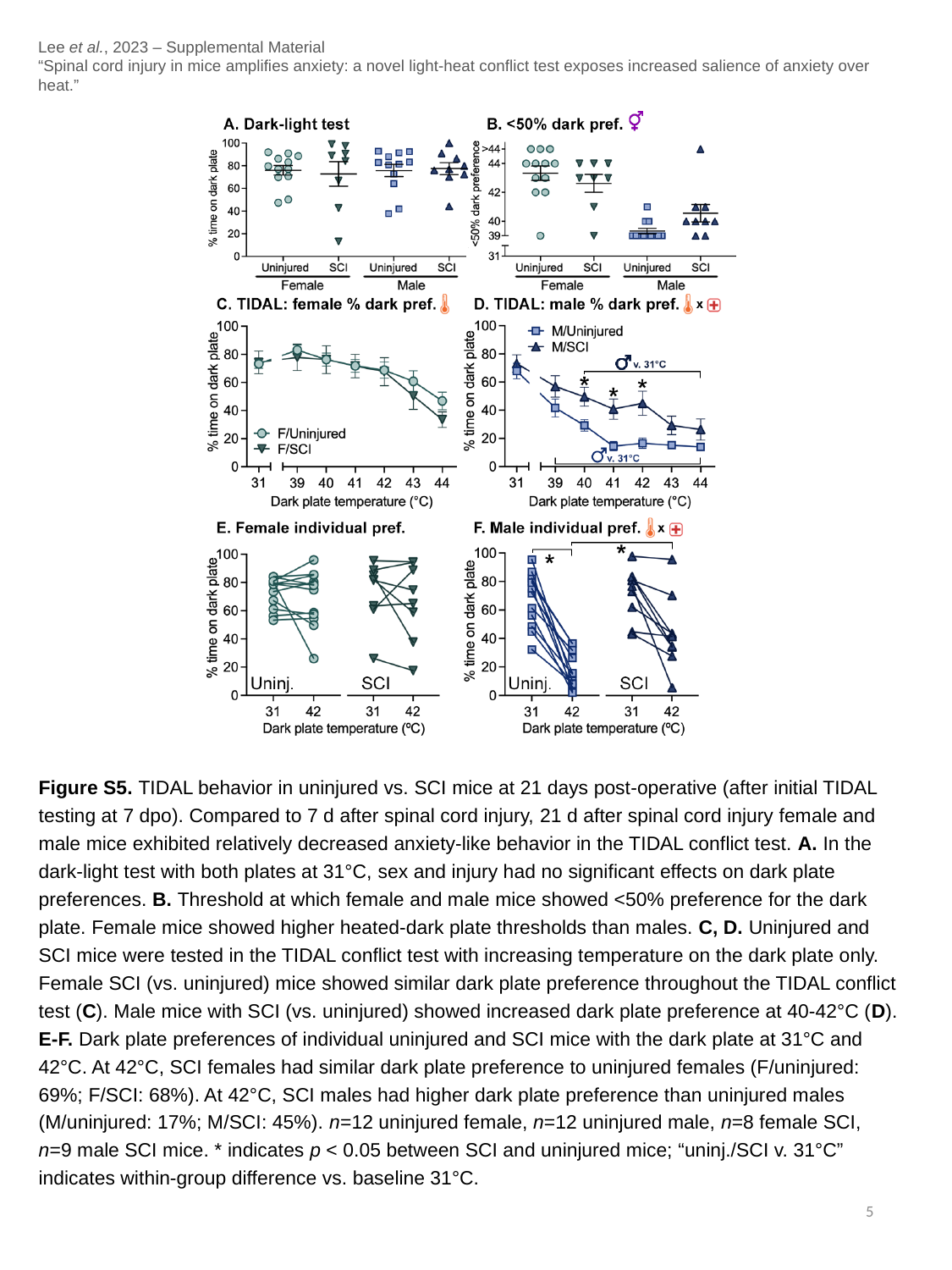

Lee et al., 2023 – Supplemental Material
“Spinal cord injury in mice amplifies anxiety: a novel light-heat conflict test exposes increased salience of anxiety over heat.”
Figure S5. TIDAL behavior in uninjured vs. SCI mice at 21 days post-operative (after initial TIDAL testing at 7 dpo). Compared to 7 d after spinal cord injury, 21 d after spinal cord injury female and male mice exhibited relatively decreased anxiety-like behavior in the TIDAL conflict test. A. In the dark-light test with both plates at 31°C, sex and injury had no significant effects on dark plate preferences. B. Threshold at which female and male mice showed <50% preference for the dark plate. Female mice showed higher heated-dark plate thresholds than males. C, D. Uninjured and SCI mice were tested in the TIDAL conflict test with increasing temperature on the dark plate only. Female SCI (vs. uninjured) mice showed similar dark plate preference throughout the TIDAL conflict test (C). Male mice with SCI (vs. uninjured) showed increased dark plate preference at 40-42°C (D). E-F. Dark plate preferences of individual uninjured and SCI mice with the dark plate at 31°C and 42°C. At 42°C, SCI females had similar dark plate preference to uninjured females (F/uninjured: 69%; F/SCI: 68%). At 42°C, SCI males had higher dark plate preference than uninjured males (M/uninjured: 17%; M/SCI: 45%). n=12 uninjured female, n=12 uninjured male, n=8 female SCI, n=9 male SCI mice. * indicates p < 0.05 between SCI and uninjured mice; “uninj./SCI v. 31°C” indicates within-group difference vs. baseline 31°C.
5
